# Shotgun Metagenomics of 361 elderly women reveals gut microbiome change in bone mass loss

**DOI:** 10.1101/679985

**Authors:** Qi Wang, Qiang Sun, Xiaoping Li, Zhefeng Wang, Haotian Zheng, Yanmei Ju, Ruijin Guo, Songlin Peng, Huijue Jia

**Affiliations:** BGI-Shenzhen, Shenzhen 518083, China; School of Future Technology, University of Chinese Academy of Sciences, Beijing, 101408, China; Department of Statistical Sciences, University of Toronto, Toronto, Canada; Department of Spine Surgery, Shenzhen People’s Hospital, Ji Nan University Second College of Medicine, 518020, Shenzhen, China; Macau University of Science and Technology, Taipa, Macau 999078, China; Shenzhen Key Laboratory of Human Commensal Microorganisms and Health Research, BGI-Shenzhen, Shenzhen 518083, China

## Abstract

Bone mass loss contributes to the risk of bone fracture in the elderly. Many factors including age, obesity, estrogen and diet, are associated with bone mass loss. Mice studies suggest that the intestinal microbiome might influence the bone mass by regulating the immune system, however there has been little evidence from human studies.

We have recruited 361 Chinese elderly women to collect data for a metagenomic-wide association study (MWAS) to investigate the influence of the gut microbiome on bone health. Gut microbiome data were produced using BGISEQ500 sequencing, BMD was calculated using Hologic dual energy X-ray machine, BMI (Body Mass Index) and age were also provided. This therefore data allows exploration of gut microbiome diversity and links to bone mass loss, as well as microbial species and modules as markers for bone mineral density. Making these data potentially useful in studying the role the gut microbiota might play in bone mass loss and offering exploration into the bone mass loss process.

## Context

Bone mass loss is a process where calcium and phosphate from the bones is reabsorbed, and instead of retaining these minerals it makes our bones weaker [1]. It is a severe and common disease in elderly population, as well as being the most common reason for fracture, giving rise to acute pain and even death [2]. Recently a new concept, “osteoimmunology”, has revealed tight interaction between the immune system and bone metabolism [3]. Interestingly, it has been widely recognized that the gut microbiota could influence host health by interacting with the host immune system [4]. But most related research [4, 5] has been carried out using mice and 16S sequencing which is of poor taxonomic resolution, low sensitivity and contains no functional related information [6]. It is crucial to explore the relationship between the gut with bone loss by shotgun sequencing. Here, we carried out the MWAS on fecal samples from 361 Chinese elderly women living in an urban setting.

## Methods

### Sampling strategy

Fecal samples were collected at the People’s Hospital of Shenzhen and immediately frozen in an 80 °C freezer for storage. The samples were then transported to BGI-Shenzhen and sequencing using the BGISEQ-500 platform (BGISEQ-500, RRID:SCR_017979), following previously published protocols [10] [11]. Quality control was performed, and adaptor and host contamination were filtered (sTab1), the high-quality non-human reads were defined as clean reads following previous methods [12] [13]. The BMD was calculated using a Hologic dual energy X-ray machine at Shenzhen people’s Hospital (sTab1). We used the T-score of BMD in the lumbar spine to represent the bone mass [14]. A sample’s T-score is a relative measure of the sample’s BMD compared to the reference population of young, healthy individuals with the same gender.

### Taxonomic and functional abundance calculation

The cleaned data were used for the annotation and profile of taxon using MetaPhlan2 (MetaPhlAn, RRID:SCR_004915)[15]. We removed species presented in less than 10% of the samples for later analysis. For functional abundance calculation, Firstly, the putative amino acid sequences were translated from the gene catalogues and aligned against the proteins/domains in the KEGG databases (release 79.0, with animal and plant genes removed) using BLASTP (v2.2.26, default parameter except that -e 0.01 -b 100 -K 1 -F T -m 8). Each protein was assigned to the KO group by the highest scoring annotated hit(s) containing at least one HSP scoring >60 bits. The relative abundance profile of KOs was determined by summing the relative abundance of genes from each KO [16] [17]. The KO abundance was as input to calculate the profile of gut metabolic modules (GMM) [18].

### Two-stage least square

Stage 1: In the first step we regress the taxonomic abundance or metabolic module abundance on the Age and BMI with linear regression and save the prediction value. The detail of the taxonomic abundance is in ‘Taxonomic abundance calculation’ part (sTab 3). The detail of the metabolic module abundance is in ‘Gut metabolic modules analysis’ part (sTab 4). This step is used to adjust the effects of Age and BMI to the contribution of BMD by the taxonomic abundance or metabolic module abundance.

Stage 2: Five-fold cross-validation is performed ten times on a random forest regression model (Y: the BMD T score; X: the prediction value from the stage 1). The error curves from ten trials of fivefold cross-validation are averaged. We chose the model which minimized the sum of the test error and its standard deviation in the averaged curve.

### Alpha-diversity and count

The within-sample diversity is calculated by profile of samples with Shannon index, as described previously [6]. Genes were considered present with more than one read map to it.

### Data Validation and quality control

The metagenomic shotgun sequencing of 361 samples was performed obtaining an average of 7.7 gigabase (Gb) clean data per sample (sTab1c). To explore the utility of this data the life and clinical index (sTab1a-b) to the T-score in our cohort was assessed and significant factors such as age and body mass index (BMI) to the microbiome were excluded, and the alterations of the gut microbiome along with the T-score were evaluated. Finally, stable regression model was built at species and module level for the cohort. The T score of the BMD in the lumbar spine was used to represent the bone mass.

### Demonstration of Utility 1. A mild gut microbiome dysbiosis seen for bone mass loss

To explore alterations of the gut along with the change of the T-score, the change in different taxonomy levels can be analyzed. Diversity at each level increases with the T-score probably caused by the increasing of pathogenic microcells in our gut. Gene (*p* = 4.53e-9, adjusted R^2^=0.0904, linear regression, sFig2b, sTab2), species (*p* = 1.17e-15, adjusted R^2^=0.162, linear regression, sFig2d, sTab2) and genus (*p* = 7.98e-14, adjusted R^2^=0.144, linear regression, Fig1b, sTab2) level are shown. In addition, the count data also shows an increase in the T-score at gene (*p* = 0.0114, adjusted R2=0.0152, liner regression, sFig2a, sTab2), species (*p* = 2.33e-5, adjusted R2=0.0465, liner regression, sFig2c, sTab2), and genus (*p* = 7.73e-10, adjusted R2=0.0992, liner regression, Fig1a, sTab2) level. Next, the top 20 abundant species are chosen (Fig1c, sTab3). The data demontrates that the *B*.*stercoris, E*.*coli, B*.*uniformis, B*.*coprocola, B*.*fragilis, E*.*rectale* and *E*.*eligens* significantly negatively associated with T score. While for the *B*.*vulgatus, B*.*massiliensis, B. caccae* and *Megamonas unclassified* display an obvious positive correlation with T score (Fig1c, sTab3). In addition, in the top 15 abundant genera, the *Eubacterium, Escherichia, Subdoligranulum, Klebsiella, Clostridium* and *Blautia* have significant negative correlation with the T-score (sFig3, sTab4). Among these genera *Eubacterium, Escherichia* are normal microorganism of the intestinal tract and can cause infection under opportunistic conditions. For positively correlated genera, the *Prevotella, Parabacteroides, Megamonas* and *Akkermansia* are included (sFig3, sTab4). For the top 10 enriched phyla, the *Bacteroidetes, Verrucomicrobia, Fusobacteria, Euryarchaeota* and *Ascomycota* are positive to BMD T-score (sFig4, sTab5), while the *Proteobaccteria, Actinobacteria, Synergistetes* and *Chlorobi* are negative (sFig4, sTab5).

**Figure 1.**
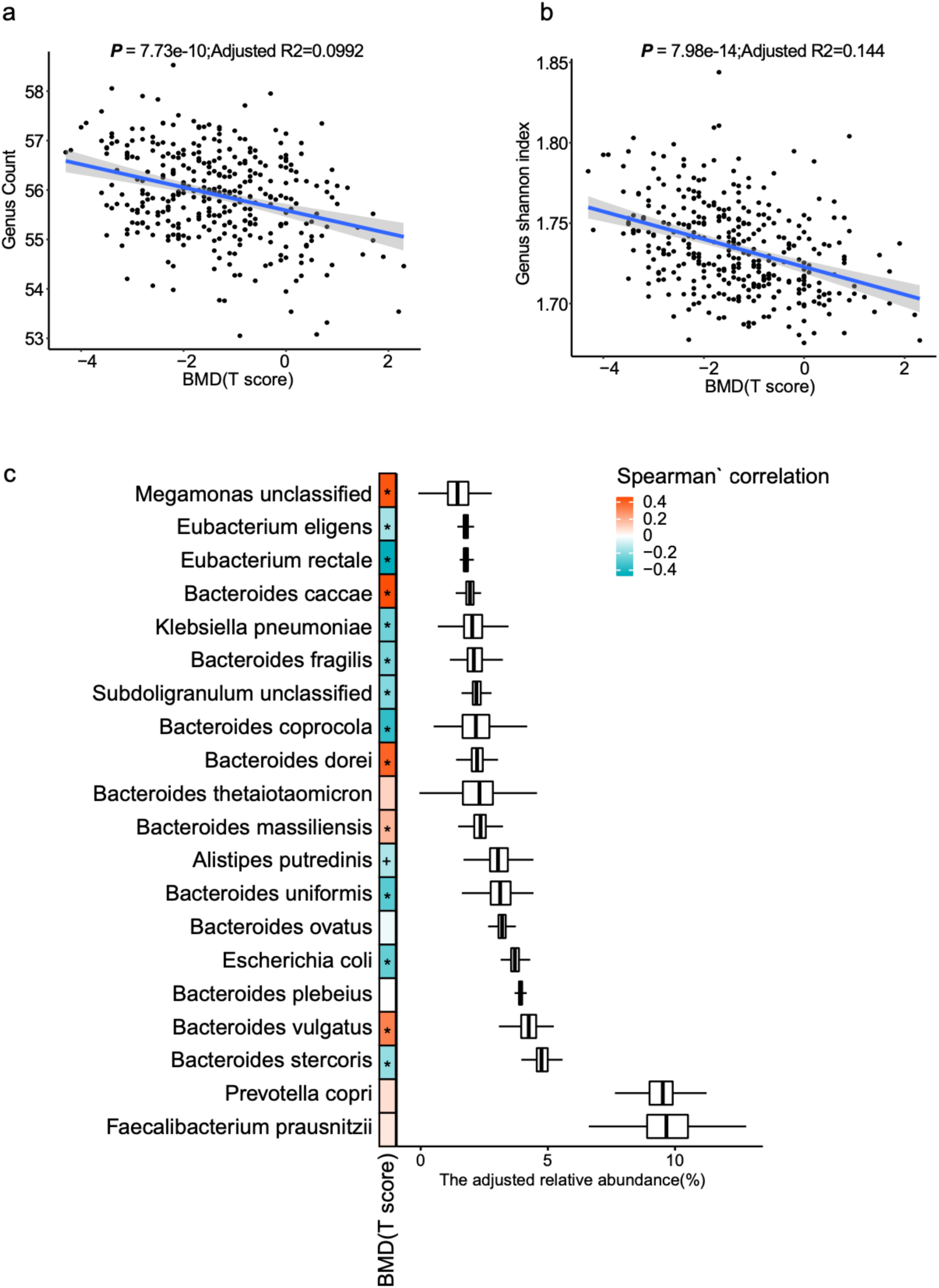
slightly increase gut microbial richness. (a-b) Richness and alpha-diversity (Shannon index) at the genus level of the two cohorts (liner regression). (c). The top 15 species. (The spearman’s correlation,’+’ for *p* < 0.05;’*’ for *p* < 0.01).

### Demonstration of Utility 2. Species linked to BMD

To select the species which have strong connection with the T-score, we use the two-stage least square method [7] to regress the species to the T-score (details are showed in methods part). The model shows a high R square (more than 0.99, sTab3a, Fig2a), and 18 species are selected. The importance of these species (Fig2b, sTab3a) are ranked in order. Spearman’ rank correlation method is used to evaluate the relationship between the selected species and the clinical indexes (Fig2c). From these findings, it is easy to see that some T-score negatively correlated species like the *Streptococcus parasanguinis, Clostridium perfringens, Haemophilus sputorum, Enterobacter aerogenes, Actinobacillus unclassified* and *Chlorobium phaeobacteroides* are negatively connected with the triglyceride (TG) levels, but positively correlate with β-Crosslaps (CROSSL) and high-density lipoprotein (HDL). As well as some T-score positively correlated species, for example the *Roseburia intestinalis* which is a butyrate-producing bacterium that could influence human’ immune system [8], *Enterobacter cloacae* and *Sutterella wadsworthensis*. These species have positive correlation to TG, but negative to CROSSL and HDL.

**Figure 2.**
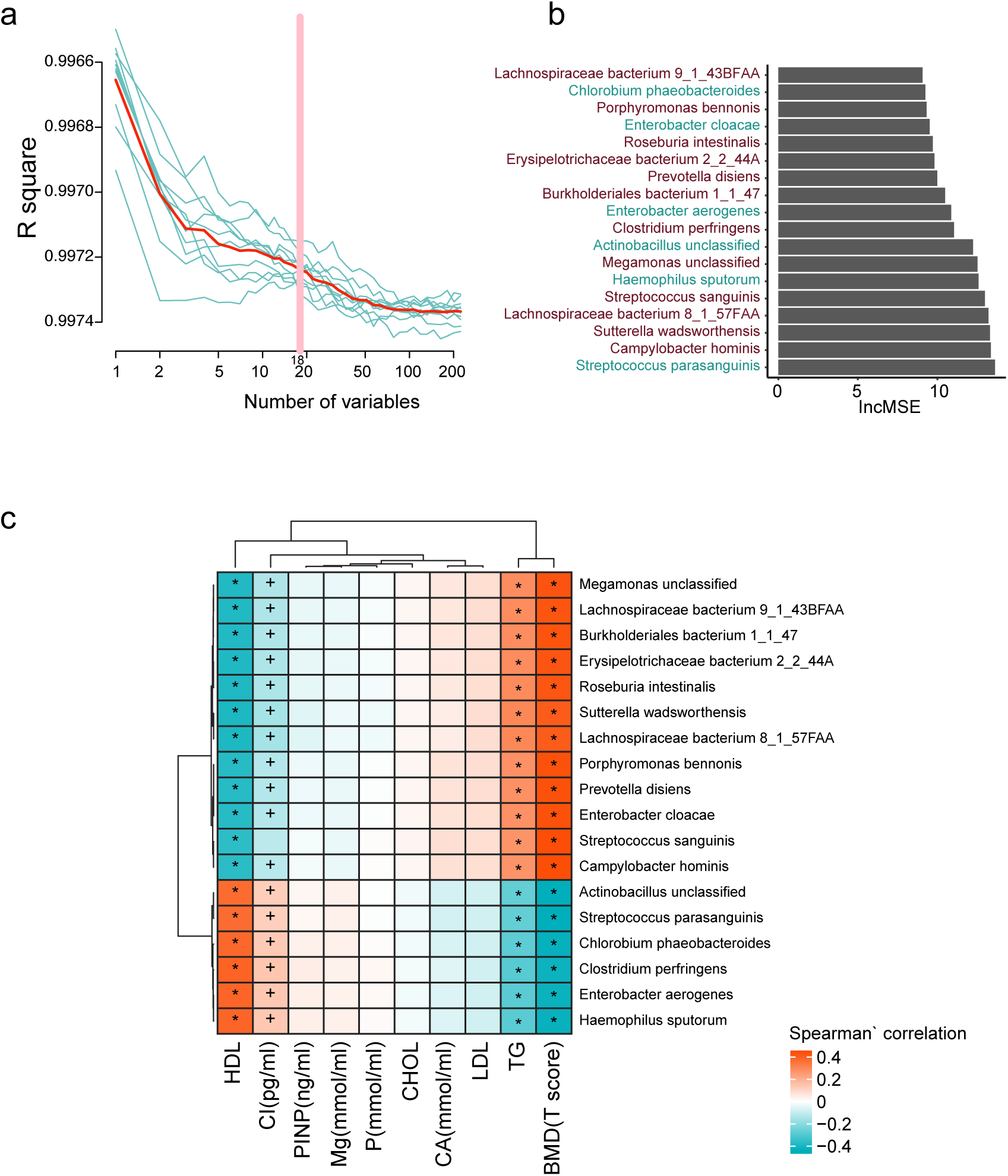
Fecal microbial species markers for BMD. (a) The R square during the Ten-time cross-validation process (the blue lines show the ten different process, the red line for the average of the ten-time cross validation, and the pink line show the best variables). (b). The lncMSE of the 18 chosen species markers. (c). The correlation between the marker with the clinical index. (Spearman’ correlation,’+’ for *p* < 0.05;’*’ for *p* < 0.01).

### Demonstration of Utility 3. Modules suggesting for bone mass loss

To find the higher correlation modules with the T-score, we also use the two-stage least square method mentioned as before. 13 modules with more than 0.99 R square (sTab4, Fig3a) are obtained by the model and plotted in rank by their importance (Fig3b, sTab4). In addition, for the correlation with clinical indexes, the negatively correlated modules like the lactate consumption, sucrose degradation, tryptophan degradation are positively associated with HDL and CROSSL, but negatively with TG. By comparison, the BMD positive modules, for example the pectin degradation, trehalose degradation, arginine degradation that can prevent bone mass loss and bone collagen breakdown in a rat model [9], mucin degradation and rhamnose degradation. These modules are positive to TG, but negative to HDL and CROSSL. For further detail on these modules please see the supplementary figures (sFig5).

**Figure 3.**
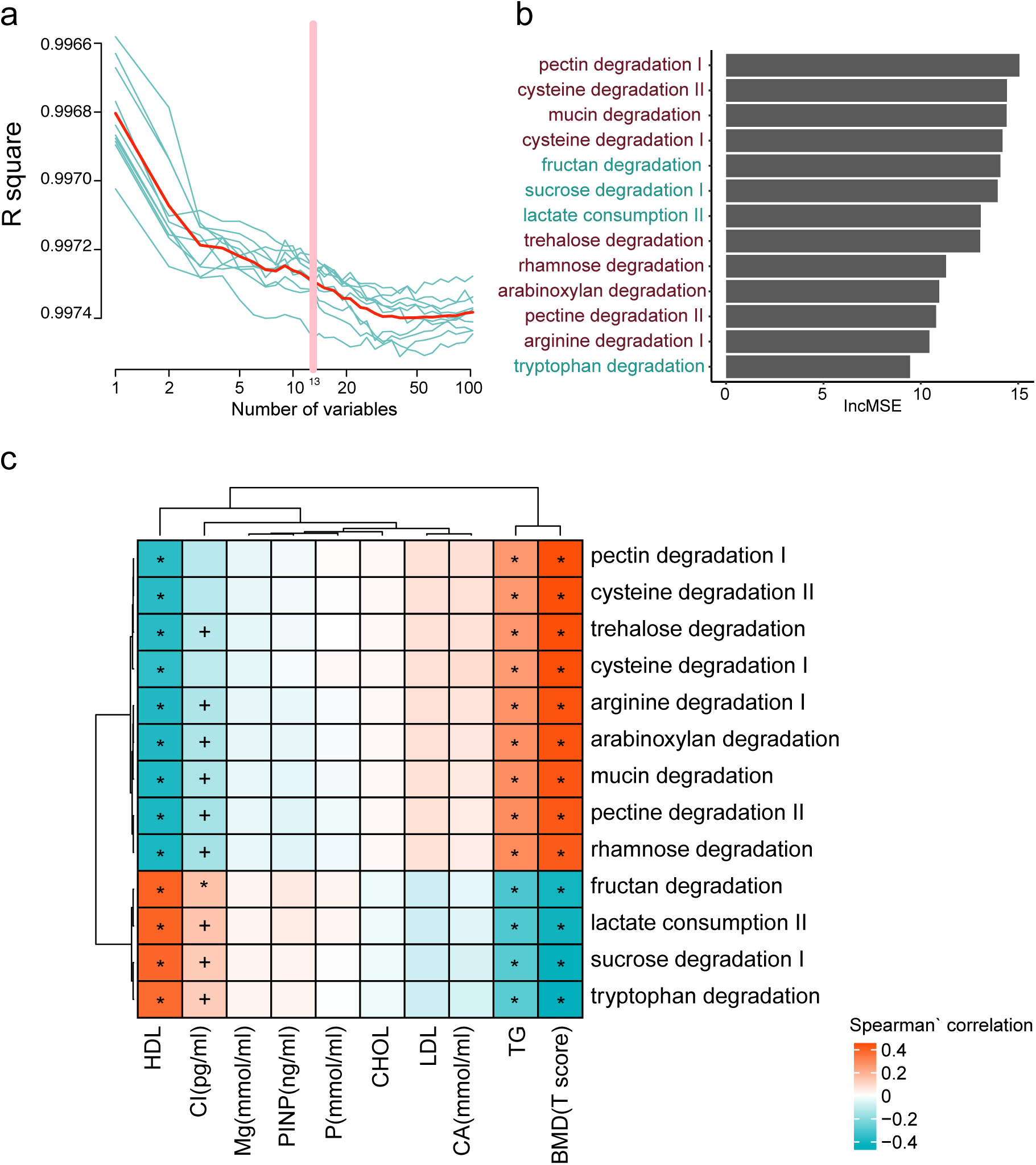
Fecal microbial modules markers for BMD. (a) The R square during the Ten-time cross-validation process (the blue lines show the ten different process, the red line for the average of the ten-time cross validation, and the pink line show the best variables). (b). The lncMSE of the 13 chosen modules markers. (c). The correlation between the marker with the clinical index. (Spearman’ correlation,’+’ for *p* < 0.05;’*’ for *p* < 0.01).

## Conclusions

We carried out the first study to explore the alteration of the gut microbiome along with bone mass loss in the 361 elderly Chinese urban women with MWAS. Firstly, taxonomy diversity is observed to increase at many levels which may be contributed by the growth of some opportunistic pathogens. In addition, some high correlated species and functional modules are also suggested which might offer us a new way for better diagnosis as well as mechanistic understandings of the bone mass loss.

Our data suggests that the gut microbiome is closely related to the process of bone mass loss in an elderly population. Although the mechanism of how do the gut microbes affect and modulate bone metabolism is not fully understood, our research indicates that gut microbiota may be novel targets for the protection of bone mass loss and provide a new avenue for the future studies and treatment of this field.

### Re-use potential

This is the first dataset where 361 high-quality metagenomics datasets were collected from elderly Southern Chinese urban women. All the clinical index also provided in the supplementary table. From initial explorations of this data we can see a slightly correlation between the metagenomics data with bone loss, and have also selected some bone loss related biomarkers in species and modules level. For re-use potential, the clinical details collected here such as the relationships of tea-drinking with bone mass or metagenomics make this a valuable dataset for further analysis. While we were unable to clearly determine any strong signals between the metagenomics and bone loss with our methods, using different statistical approaches others may find novel insight in this data. And with the still moderate sample size and detailed information on a number of clinical features, it may be a very useful dataset to combine and compare to other metagenomics datasets.

## Supporting information

sFig1

sFig2

sFig3

sFig4

sFig5

Stab1

sTab2

sTab3

sTab4

sTab5

sTab6

## Ethics approval and consent to participate

This study was approved by the Institutional Review Board on Bioethics and Biosafety at Shenzhen people’s Hospital (LL-KY-2019506) and BGI (BGI-IRB 19126).

### Availability of data and materials

The sequencing reads from each sequencing library have been deposited at EBI with the accession number: PRJNA530339 and the China National Genebank (CNGB), accession number CNP0000398.

## Abbreviations

BMD: Bone Mineral Density
BMI: body mass index
Gb: gigabase
MWAS: metagenomic-wide association study
GMM: gut metabolic module
TG: triglyceride
HDL: high-density lipoprotein
CROSSL: β-Crosslaps

