## Supplementary figures and images for "Shotgun Metagenomics of 361 elderly women reveals gut microbiome change in bone mass loss"

### sFig1

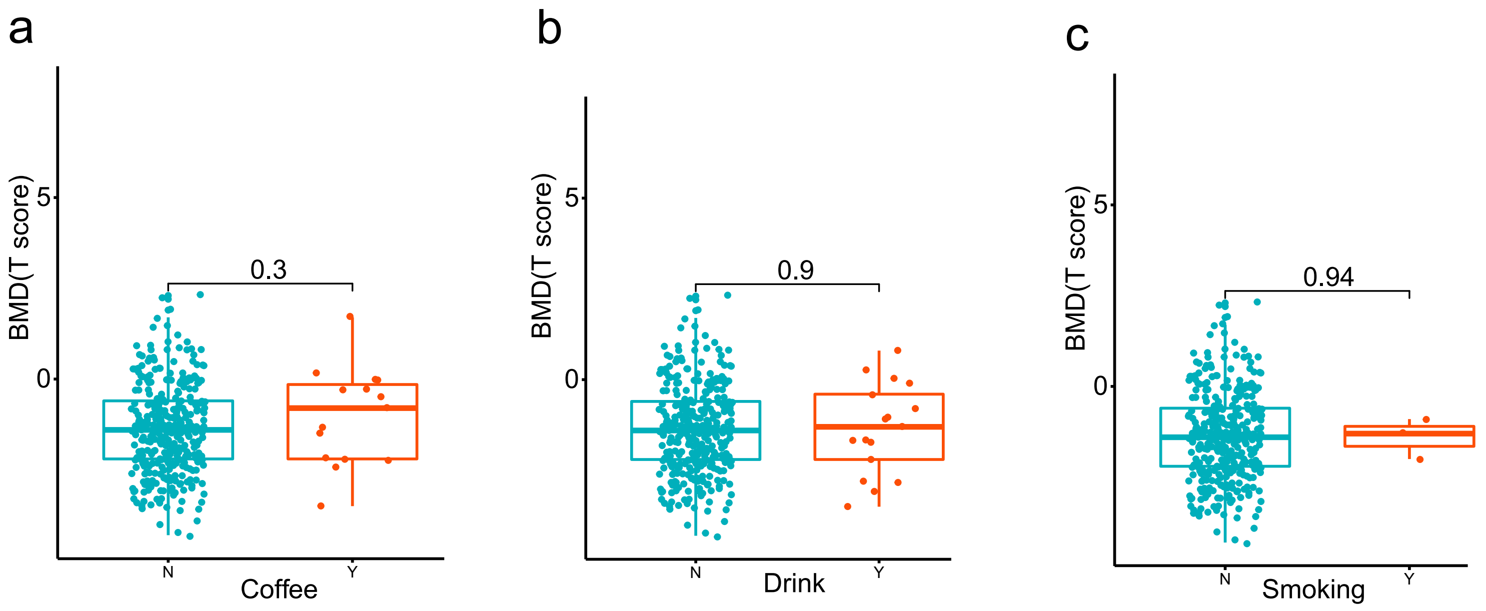

### sFig2

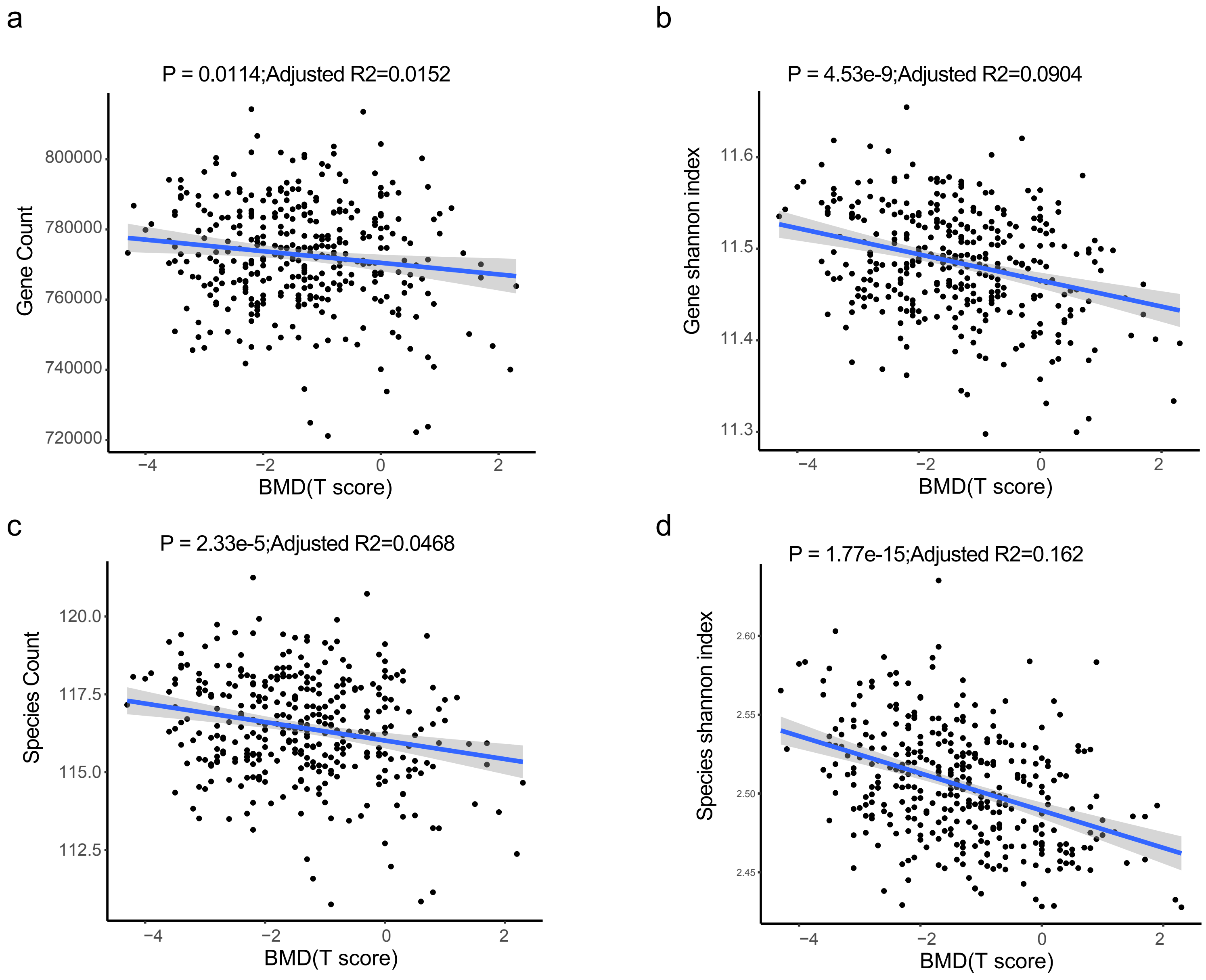

### sFig3

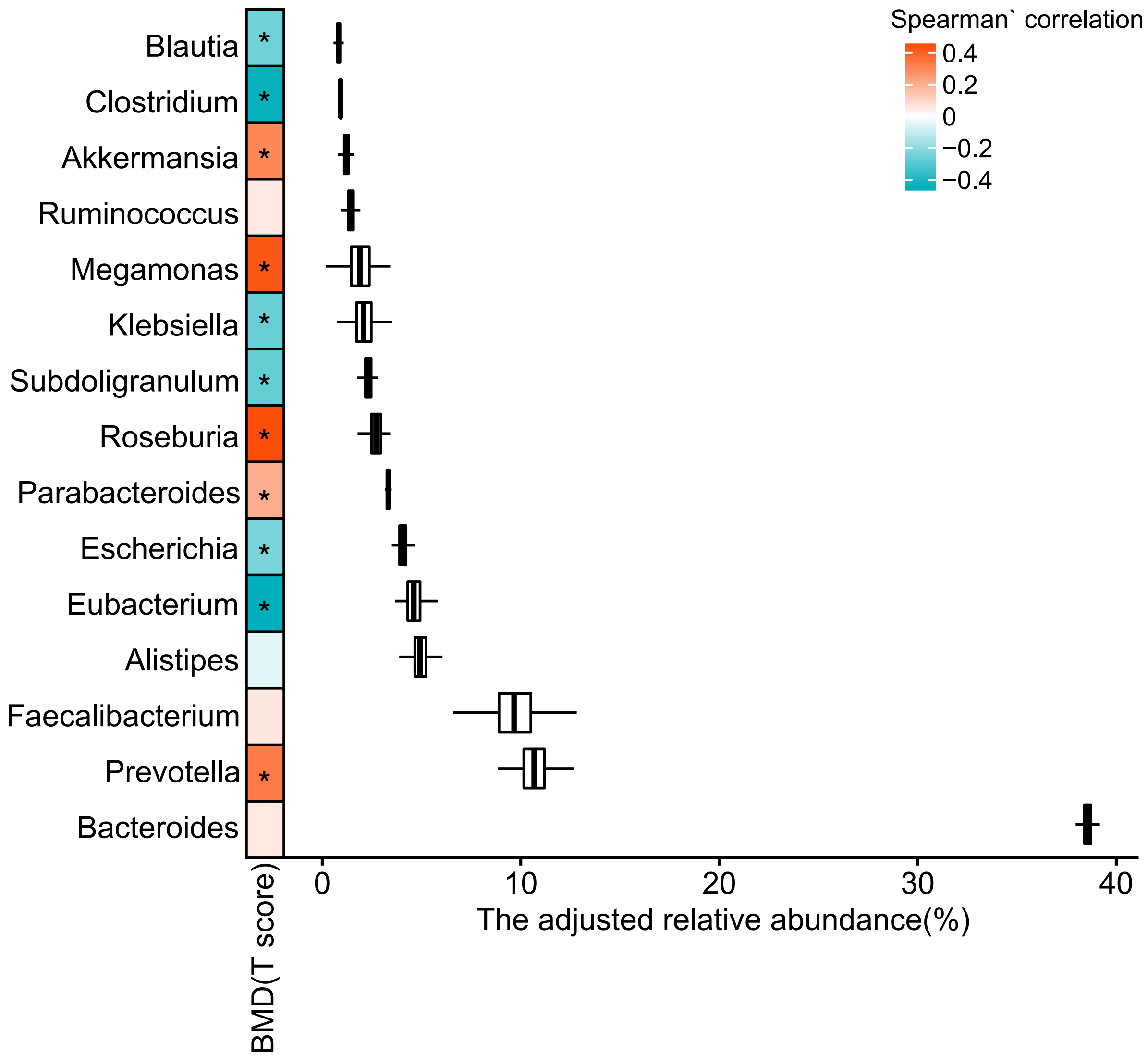

### sFig4

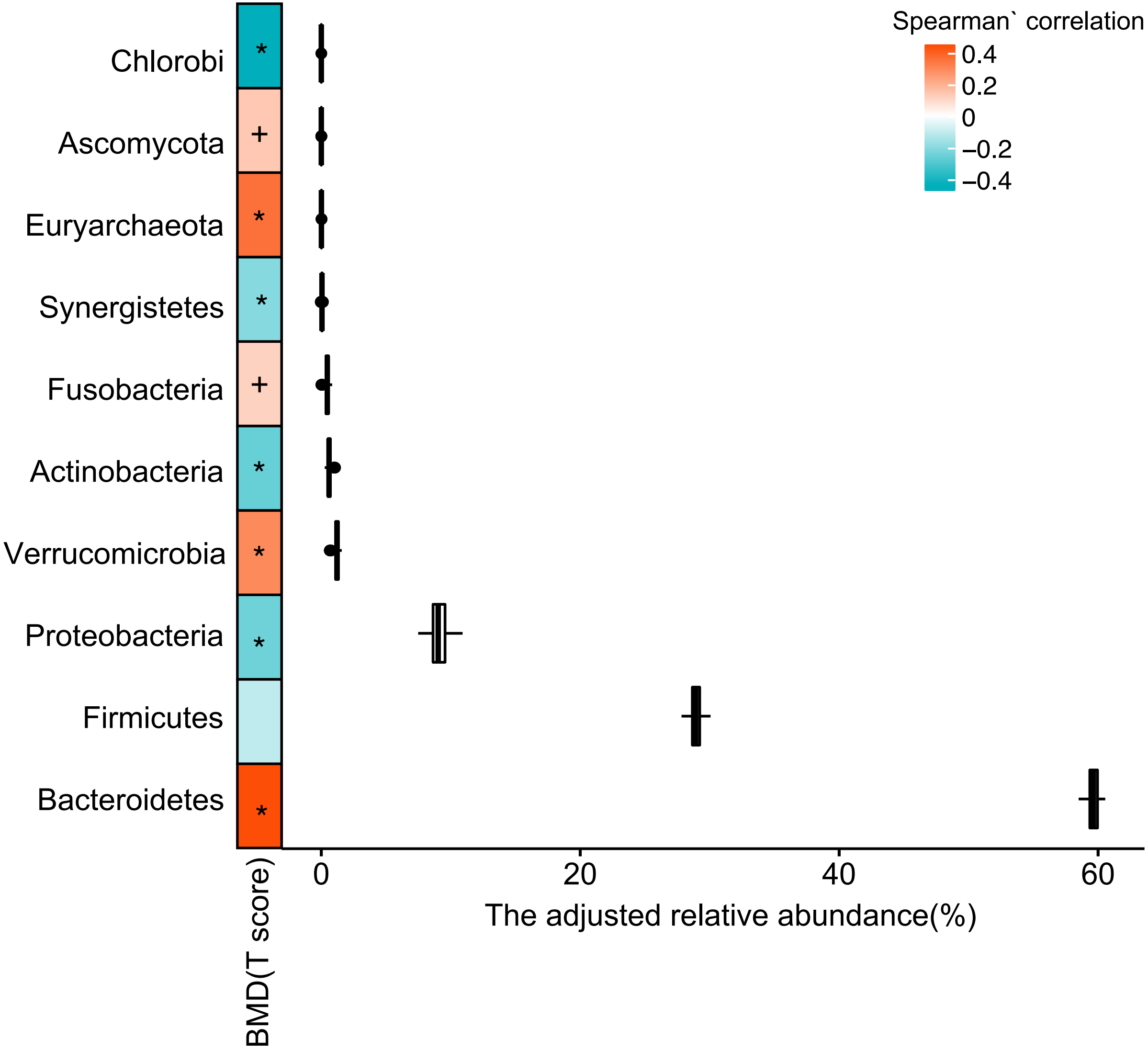

### sFig5

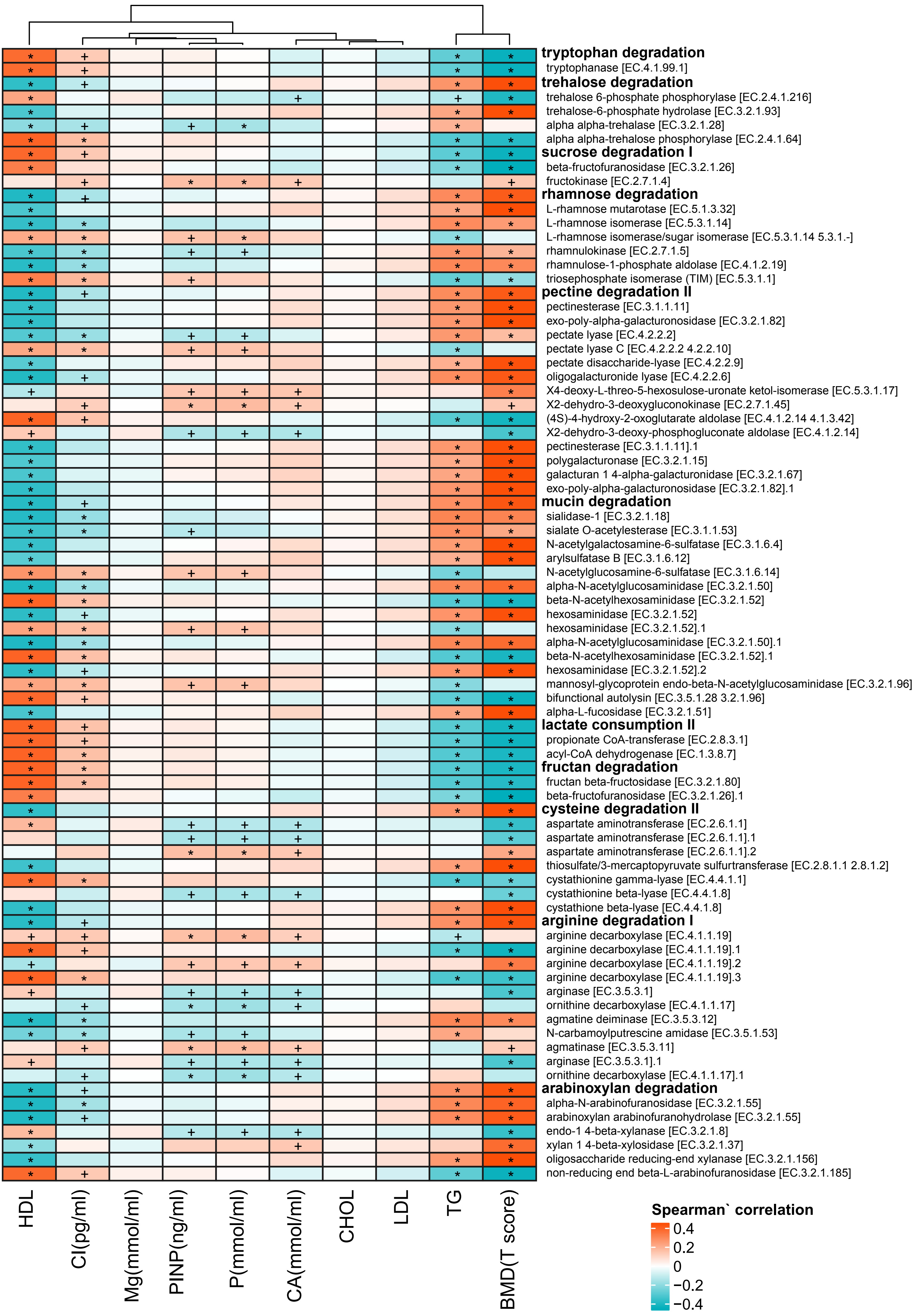
